# Evidence of inbreeding depression on stature in Brown Swiss cattle

**DOI:** 10.1101/2025.01.22.634247

**Authors:** Qiongyu He, Jessica Deuber, Franz R. Seefried, Hubert Pausch, Naveen Kadri

## Abstract

**Background:** Small effective population size and the disproportionately large use of few genetically superior bulls in artificial insemination lead to extensive runs of homozygosity and an increased risk of homozygosity for deleterious alleles in domestic cattle, which may cause inbreeding depression. The adverse effects of inbreeding on phenotypic performance are well established, but the genetic variants contributing to inbreeding depression remain largely unknown. This study aimed to analyse the impacts of inbreeding on stature (measured as height at the sacral bone) in a cohort of 15,306 Brown Swiss (BS) cows that have imputed genotypes at 20 million sequence variants and stature measurements as height at the sacral bone.

**Results:** The average genomic inbreeding coefficient of the 15,306 BS cows estimated from runs of homozygosity (ROH) was 0.369 (± 0.022). We found a loss in stature, decreasing from a height of 0.076 cm at the sacral bone per 1% increase in inbreeding (p = 1.94e-09). Contributions to inbreeding depression were significant for long (> 2 Mb), medium (0.1 – 2 Mb), and short (50 kb – 0.1 Mb) ROH (p = 1.29e-12, p = 3.20e-04 and p = 1.77e-06, respectively), suggesting that both ancient and recent inbreeding have negative effects on stature. Non-additive association testing identified a novel recessive quantitative trait locus (QTL) for stature on chromosome 25 with the most significantly (p = 2.35e-21) associated SNP residing at 14,535,327 bp. Cows homozygous for the alternate allele of the top associated SNP were 2 cm shorter than heterozygous and reference allele homozygotes. Fine mapping of the QTL identified a splice donor variant (rs447836030 at 25:14,515,474) of the gene *ABCC6* encoding ATP binding cassette subfamily C member 6 which causes exon skipping as both a positional and functional candidate causal variant.

**Conclusions:** Our study reveals evidence for inbreeding depression on stature in a large cohort of BS cattle. We also uncover a recessive QTL that decreases stature through non-additive association testing. This QTL harbors a high-impact variant affecting a splice donor site of *ABCC6* which leads to exon skipping, thereby possibly contributing to inbreeding depression. Accumulating non-lethal deleterious alleles in ROH may reduce the overall fitness of the BS cattle population.

## Background

Inbreeding refers to matings between individuals who share a common ancestry. Inbreeding occurs frequently in species with a small effective population size and a disproportionate contribution of few individuals to the next generation (1). A direct consequence of inbreeding is an increased probability of homozygosity for recessive deleterious alleles (1), resulting in reduced fitness and referred to as inbreeding depression.

The inbreeding coefficient (F) quantifies the level of inbreeding (2). Traditionally, this measurement relies on pedigree information (F_PED_) to estimate the proportion of alleles that are identical by descent (IBD) (3,4). Depth, accuracy, and completeness of the pedigree impact the accuracy of F_PED_ (5). Pedigree records cannot accurately describe variation in Mendelian sampling and linkage during gamete formation among offspring from the same mating type (4,6,7). Thus, F_PED,_ can differ substantially from the realized IBD, which can be estimated using genomic data.

Several methods exist to estimate inbreeding from genomic data, such as the correlation of uniting gametes (F_UNI_), genomic relationship (F_GRM_), excesses homozygosity (F_HOM_) or runs of homozygosity (ROH) in the autosomal genome (F_ROH_), among others (8–13,5). However, the optimal approach for precise measurement of genomic inbreeding has yet to be established (13,14). Genomic inbreeding quantified through ROH (F_ROH_) enables assessing IBD sharing at the chromosome and whole-genome levels (5,12–14) more accurately than using pedigrees (6,15,16).

The age of inbreeding can be estimated from the length of ROH (12), where longer ROH are often the result of more recent inbreeding events. ROHs are generally enriched for homozygous deleterious variants (17,18), though the degree of enrichment across length categories varies between species due to different selective pressures and demographic histories (17–22). Long ROH are enriched for deleterious mutations in species like humans, likely because purifying selection has had less time to purge them (23); empirical evidence from cattle presents a contrasting result, showing that shorter ROH are more strongly enriched for deleterious alleles (18). The accumulation of deleterious mutations in the homozygous state can result in decreased fitness, possibly culminating in inbreeding depression at the population level (24).

Inbreeding depression on fitness traits has been established in various cattle breeds (21,25) but the underpinning genetic and molecular causes are largely unknown (26). Genome-wide association studies (GWAS) have provided only limited insights into the genetic architecture of inbreeding depression (21,26,27), mainly because they typically assume additive inheritance. A few non-additive GWAS have identified recessive loci impacting fitness in cattle (28–30), suggesting that association testing with non-additive models might facilitate characterizing the genetic architecture of inbreeding depression.

This study examines how inbreeding impacts stature in Brown Swiss (BS) cattle and assesses how both recent and ancient ROH contribute to inbreeding depression. We also conduct non-additive association testing to investigate the genetic architecture of inbreeding depression on stature. Further, we examine the distribution of deleterious allele homozygosity within ROH to gain insights into how inbreeding could affect fitness in general.

## Methods

### Genotypes

Genotypes for 61,250 cattle from the Original Braunvieh (OB) and Brown Swiss (BS) breeds and crosses between the two breeds were obtained using 10 SNP arrays comprising between 20k and 777k SNPs. The positions of the SNPs corresponded to the ARS-UCD1.2 (bosTau9) (31) assembly of the bovine genome. Only autosomal SNPs were considered in our analyses. We employed PLINK (version 2.0) (32) for quality control separately for each subset of cattle genotyped with the same array. Samples and SNPs with more than 10% missing genotypes and SNPs deviating significantly (p < 10e-08) from Hardy-Weinberg proportions were filtered out.

Sequence variant genotypes were imputed into the array-typed genotypes with a two-step imputation procedure using Beagle (version 5.2) (33). In the first step, genotypes of 682,946 autosomal SNPs typed with the Illumina Bovine HD SNP Bead Chip were imputed using a reference panel comprising 1,183 BS and OB samples. In a second step, genotypes for 36,559,070 sequence variants were imputed into the previously imputed high-density genotypes using a multi-breed reference panel of 658 sequenced samples, including 298 BS and OB animals. The procedures employed for sequencing, alignment and variant calling of the sequenced reference panel have been described earlier (28).

### Principal component analysis and breed assignment

A genomic relationship matrix (GRM) was built from 16,289,812 autosomal SNPs with minor allele frequencies (MAF) greater than 0.01 using the “--grm” flag of the GCTA (version 1.93.2a) software (34). The principal components (PC) of the GRM were obtained with the “--pca” flag of GCTA. The first two PCs were used to assign breeds (Additional file 1, Supplementary Figure S1). This procedure identified 50,293 BS (0.000 < PC1 < 0.004, –0.010 < PC2 < 0.020) animals.

### Phenotypes

Stature was measured as height (in cm) at the sacral bone in 15,306 genotyped BS cows. All considered measurements were recorded in the first lactation. We fitted the stature measurements using a linear mixed model implemented in R (35) to account for non-genetic factors. The fitted model included age and days since calving as fixed effects, and expert and farm as random effects. The residuals from this model were used as phenotypes for GWAS and heritability estimation.

### Heritability estimation

Additive and dominance genomic relationship matrices (GRM) were constructed using 14,536,150 autosomal SNPs that had MAF > 0.01 in 15,306 BS cows using GCTA with the “--make-grm” and “--make-grm-d” flags, respectively. Both GRMs were fitted simultaneously using the “--mgrm” flag to estimate genome-wide additive (h^2^) and dominance (8^2^) heritability with a REML (“--reml”) algorithm implemented in GCTA. We also included the first four PCs of the additive GRM to account for hidden relationships. The statistical significance of the 8^2^ was determined using a log-likelihood test implemented in GCTA.

We partitioned h^2^ and 8^2^ across the 29 autosomes. Additive and dominance GRMs were constructed for each chromosome separately, considering SNPs with MAF > 0.01 using the “--make-grm” and “--make-grm-d” flags of GCTA, respectively. To estimate the contribution of each chromosome to h^2^, we fitted all 29 additive GRMs simultaneously using the “--mgrm” flag while accounting for the top four PCs of the additive GRM. For partitioning 8^2^, we included the additive GRM derived from genome-wide SNPs and 29 dominance GRMs simultaneously. Heritability was set to 0 when the estimate was < Vp/10e-08, where Vp is the phenotype variance. We used one-tailed t-tests to evaluate the significance of each chromosome’s contribution.

### Genome-wide association study (GWAS)

The association between stature and imputed sequence variants was tested assuming additive and non-additive inheritance using a mixed linear model implemented in GCTA (34). The additive GRM and the top four PCs of the additive GRM were included as covariates to account for relatedness and population stratification. Additive association testing was conducted using GCTA’s default option with the “--mlma” flag that estimates the allele substitution effect (b). For the non-additive association testing, we estimated the dominance effect

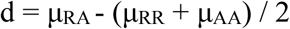

where μ is the phenotypic means among the three genotype classes for the reference (R) and alternative (A) allele following the genotype coding suggested by Zhu et al. (37). Genotypes were coded as RR = 0, RA = 2p and AA = 4p−2, where p is the alternate allele frequency estimated in the GWAS cohort of 15,306 BS animals (37). SNPs with p values below 5e-08 were considered genome-wide significant.

### Fine mapping

The Variant Effect Predictor software (VEP) (38) was used to predict functional consequences of sequence variants based on the Ensembl (version 104) annotation of the bovine reference genome. Linkage disequilibrium (LD; r^2^) between the top trait-associated SNP and all other SNPs within a 2-Mb window was calculated using PLINK. All protein-coding variants in high LD (r^2^ > 0.8) with the top trait-associated variant were considered as positional candidate causal variants.

### Functional characterization of a candidate causal variant

Testis transcriptomic data from a previously established cohort of 117 deep RNA and DNA sequenced BS bulls (39) were investigated to study impacts of a splice donor variant (rs447836030) in *ABCC6* on gene expression and splicing. Gene-level expression (in transcripts per million (TPM)) was estimated with the QTLtools quan function (40) as previously described (39) and regressed on the genotypes of the rs447836030 variant. RNA coverage was averaged over the annotated exons of *ABCC6* (transcript: ENSBTAT00000050284) using mosdepth (v0.3.6) (41) and standardised per sample based on the number of properly paired reads that were mapped to the bovine reference sequence using STAR (version 2.7.9a) (42). Differential splicing was analysed using Leafcutter (version 0.2.9) (43) as previously described (39).

### Runs of homozygosity and genomic inbreeding

Runs of homozygosity (ROH) in the imputed sequence variant genotypes of 15,306 cows were identified using PLINK. We used sliding windows of 50 SNPs, allowing for one heterozygous SNP per window and requiring a minimum number of 100 SNPs within a ROH. We set the minimal acceptable length of ROH to 20 kb (44,45) while keeping the rest of the parameters at their default values.

Genomic inbreeding coefficients (F_ROH_) were computed with the R package ‘detectRUNS’ (46) as L_ROH_ / L_ats_, where L_ROH_ is the total length of ROH in an individual’s genome, and L_ats_ is the total length of autosomes. We categorized ROH into three groups: short (50 kb – 0.1 Mb), medium (0.1 Mb – 2 Mb), and long (> 2 Mb) (47) and estimated the corresponding genomic inbreeding as F_long_, F_medium_, and F_short_. We used the formula g = 100 / (2 x L), where L is the average length of ROH to estimate the average number of generations to the most recent ancestor (20).

### Inbreeding depression

We regressed stature on F_ROH_ using a linear mixed model accounting for pedigree relationships using AIREMLF90 (48) to investigate possible adverse effects of inbreeding (hereafter referred to as inbreeding; ID). The model also included four PCs of the genomic relationship matrix, age, and “days since calving” as fixed effects, and expert and farm effects as random effects. The same model that simultaneously fitted F_long_, F_medium_, and F_short_ was used to estimate the contributions of short, medium, and long ROH to ID. The significance of ID was tested using Student’s t-test in R (49).

### Distribution of functional variants in ROH

We used the SIFT score integrated within VEP to predict the functional impact of variants segregating in the BS population. Variants identified as high-confidence deleterious mutations (n = 15,708) and high-confidence tolerant (non-deleterious) mutations (n = 40,439) were retained to study the distribution of functional variants in ROH (18).

We adopted equations 10 and 13 from Szpiech et al. following the same notations (17), referred to here as M1 and M2, to investigate the distribution of homozygosity for deleterious and non-deleterious alleles within and outside ROH.

Briefly, the model

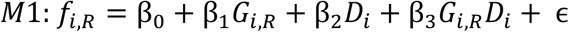

was fit to compare the enrichment of homozygous deleterious alleles and homozygous non-deleterious alleles within ROH. Where *f_i,R_* is a vector of 30,612 observations on the proportion of homozygous deleterious (d) alleles(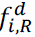) (n = 15,306) and the proportion of homozygous non-deleterious (n) alleles (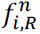) (n = 15,306) within any size of ROH of the i-th individual? *G_i,R_* is the genomic inbreeding coefficient or the proportion genome in any ROH (F_ROH_) for the i-th individual. *D_i_* is the indicator variable coded as 0 when the observation is for a non-deleterious allele (baseline) and 1 when it is for a deleterious allele. One-tailed t-tests on β_2_ (intercept) and β_3_ (slope) were performed to evaluate whether the intercept and slope for the proportion of deleterious allele homozygosity were higher than that for the proportion of non-deleterious allele homozygosity within ROH.

The same model was used to investigate the distribution of homozygosity of deleterious alleles within ROH of different ROH size category by considering the proportion of deleterious allele homozygosity within ROH of a size category and the corresponding proportion of an individual’s genome in that ROH size category.

A second model

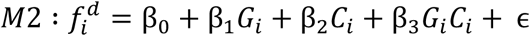

was fit to compare the enrichment of homozygous deleterious alleles within ROH of different size categories. All possible comparisons, namely, between short vs medium, short vs long and medium vs long, ROH were carried out considering the longer of the two as the baseline case. Where 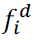 is a vector of 30,612 observations on the proportion of homozygosity for deleterious alleles in the two ROH categories compared: long (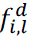), medium (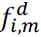) and short (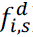). *G_i_* corresponds to genomic inbreeding coefficient corresponding to the ROH categories being compared for the individual *i*. *C_i_* is the indicator variable representing the ROH categories, where the observations for the longer ROH categories are coded as 0 (baseline), and those from shorter ROH are coded as 1. One-tailed t-tests on β_2_ (intercept) and β_3_ (slope) were performed to evaluate whether the intercept and slope for shorter ROH were significantly higher than those for longer ROH.

## Results

### Genomic inbreeding in Brown Swiss cattle

We estimated the proportion of an individual’s genome in runs of homozygosity (F_ROH_) to assess genomic inbreeding in 15,306 BS cows. An average number of 4,003 (± 109) ROH per individual that were 186 kb (± 17 kb) long was identified across the 29 autosomes, resulting in F_ROH_ between 0.276 and 0.542, with a mean of 0.369 (± 0.022). However, F_ROH_ varied substantially along the chromosomes (Figure 1A). The lowest mean F_ROH_ value (0.217) was observed for BTA23, likely due to an increased genetic diversity within the major histocompatibility complex region (50). The largest mean F_ROH_ value (0.399) was observed for BTA6, which contains numerous quantitative trait loci (QTL) associated with economically relevant traits that are likely under selection in BS cattle (Fig. 1A) (51–54).

**Figure 1.**
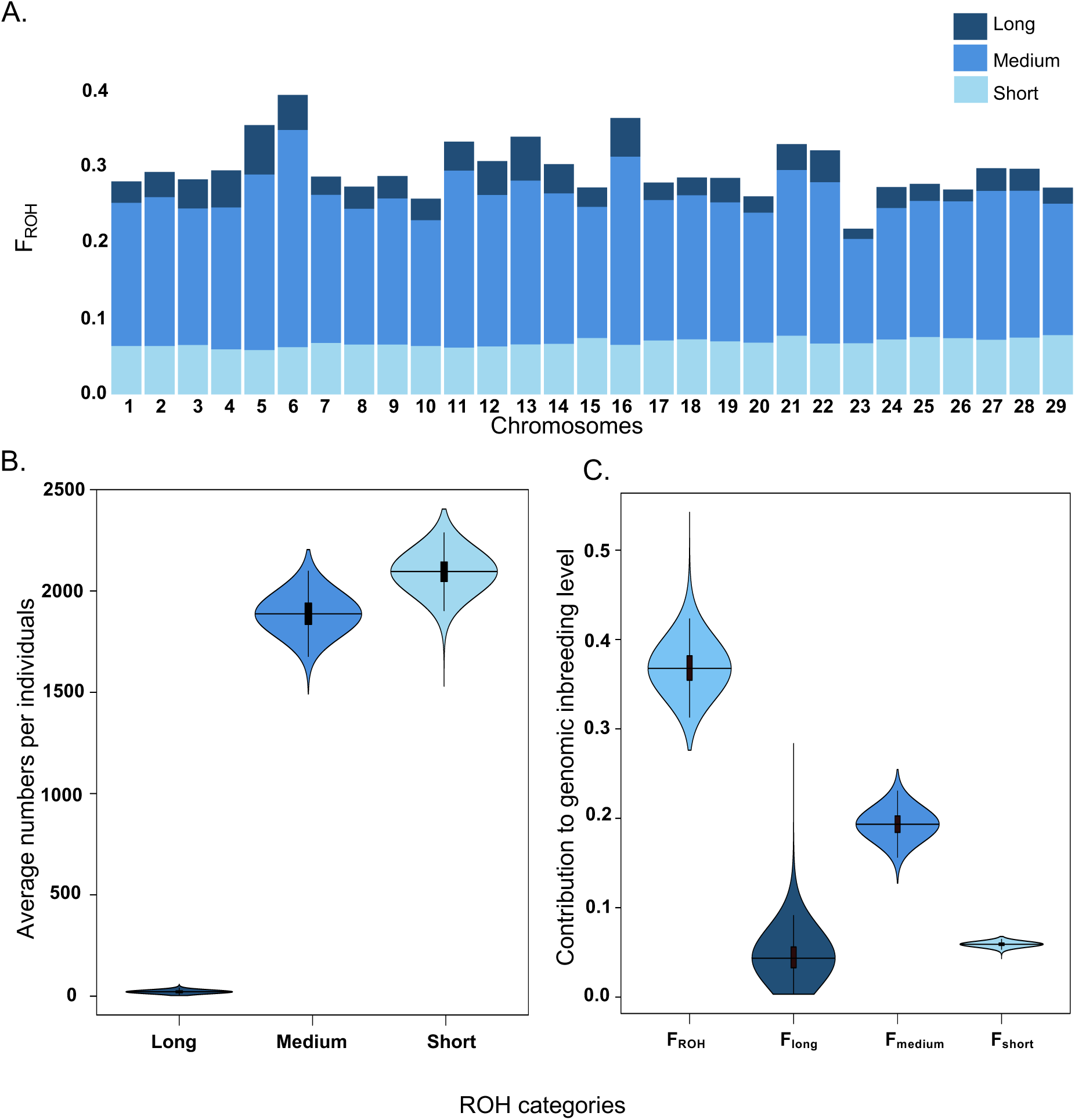
Genomic inbreeding in 15,306 Brown Swiss cattle. **A.** Proportion of autosomes in short, medium and long ROH. **B**. Average numbers of short, medium, and long ROH detected per individual. **C.** Contribution of different types of ROH to the genomic inbreeding.

We quantified the proportion of an individual’s genome in short (50 kb – 0.1 Mb), medium (100 kb – 2 Mb) and long ROH (> 2 Mb) as F_short_, F_medium_ and F_long_, respectively, to investigate ancient and recent inbreeding. Short ROH were the most frequent category (mean count: 2094 ± 74; mean length: 70 ± 0.3 kb), accounting for 52.30% of all ROH identified. Medium-sized ROH (mean count: 1888 ± 79; mean length: 255 ± 11 kb) were nearly as prevalent as short ROH and accounted for 47.15% of all identified ROH, whereas long ROH (mean count: 22 ± 6; mean length: 5205 ± 1382 kb) were rarer, accounting for only 0.55% of all ROH (Fig. 1B).

Medium-sized ROH contributed the most to the overall inbreeding (mean F_medium_: 0.193 ± 0.014) and long ROH contributed the least (mean F_long_: 0.046 ± 0.019) (Fig. 1C). Both these estimates were positively correlated with the overall F_ROH_ (r = 0.66 and 0.82 respectively, p < 2.2e-308), while F_short_ with intermediate contribution (mean F_short_ = 0.059 ± 0.002) was negatively correlated with F_ROH_ (r = –0.65; p < 2.2e-308). Short ROH reflect ancient inbreeding which occurred approximately 714 generations ago, and medium and long ROH reflect more recent inbreeding which occurred approximately 196 and 10 generations ago, respectively.

### Inbreeding has adverse effects on stature

To investigate potential effects of inbreeding on complex phenotypes, we considered stature as a prototypical trait. A linear mixed model was employed to regress individual cow stature on the inbreeding coefficient (F_ROH_). The stature of cows measured as height at the sacral bone was between 127 cm and 163 cm (mean = 147.91 ± 3.89 cm). A significant inverse association was found between F_ROH_ and stature (beta = –7.56, p = 1.94e-09). This suggests that for every 1% increase in F_ROH_, stature at the sacral bone decreased by 0.076 cm which is equivalent to a 1.9 cm stature reduction in offspring from half-sib matings (compared to offspring from matings between unrelated individuals).

A linear mixed model which fitted F_long_, F_medium_, and F_short_ simultaneously provided evidence that all three categories of ROH contributed to ID. The most significant negative effect was observed for F_long_ (beta = –13.196, p = 1.29e-12) followed by F_short_ (beta = –85.231, p = 1.77e-06), and F_medium_ (beta = –8.121, p = 3.20e-04).

### BTA25 contributes disproportionately to the dominance heritability of stature

We constructed dominance and additive genomic relationship matrices to partition the genetic variation of stature into non-additive and additive components. This analysis revealed a high h^2^ of 0.5347 (± 0.0113) and a low yet significant 8^2^ of 0.0348 (± 0.0067, p = 3.94e-10) for stature. We partitioned the h^2^ and 8^2^ to the 29 bovine autosomes to quantify individual chromosomal contributions to genetic variation. The contributions of all autosomes to the h^2^ of stature were significantly different from zero (Fig. 2A). However, only BTA25 (0.0045 ± 0.0014; p = 5.1e-04), BTA26 (0.0032 ± 0.0015; p = 0.014), BTA7 (0.0038 ± 0.0019; p = 0.023) and BTA29 (0.0027 ± 0.0014; p = 0.026) had contributions to the 8^2^ that were significantly different from zero (Fig. 2B). Length of autosomes was positively correlated with h^2^ (ρ = 0.47, p = 0.01) (Fig. 2A) but was not correlated (p = 0.22) with 8^2^ (Fig. 2B). BTA25 was an outlier, as it showed a disproportionately high contribution to both h^2^ and 8^2^. Variants on BTA25 explained 12.80 % and 7.65 % of the genome-wide δ^2^ and h^2^, respectively, despite BTA25 being the shortest of all autosomes, i.e., it contains only 1.7% of the autosomal sequence.

**Figure 2.**
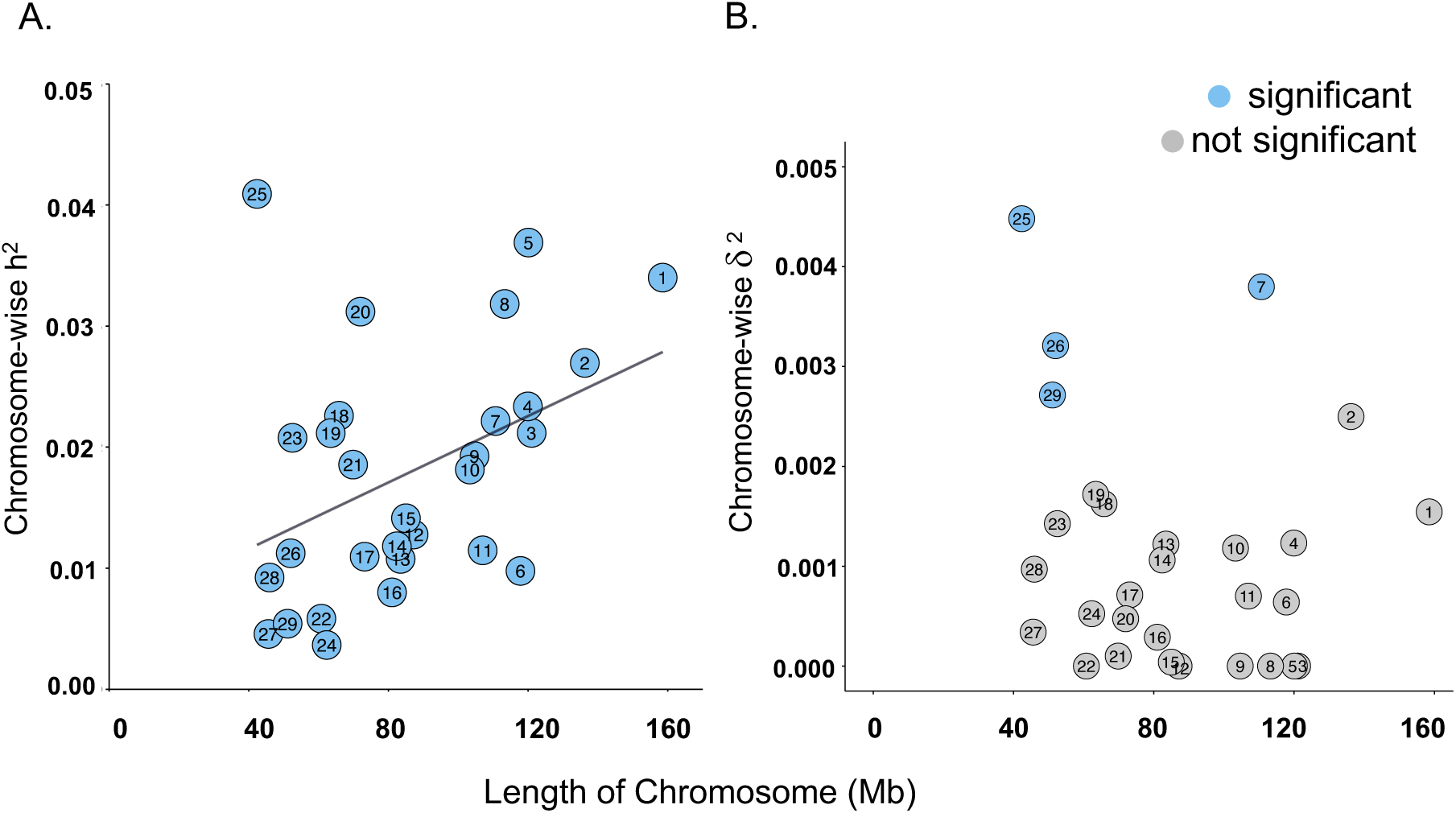
Autosomal partitioning of the heritability for stature. Contribution of autosomes to additive (**A**) and dominance (**B**) heritability of stature in Brown Swiss cattle. Statistically significant (p < 0.05) estimates are plotted in blue. The chromosomes are ordered by their size along the x-axis. The black line in **A** shows the correlation of chromosome-wise heritability and chromosome length.

### Non-additive GWAS identifies a stature-associated recessive variant

To investigate a potential contribution of large-effect loci to ID on stature, we performed genome-wide association studies between stature and imputed sequence variant genotypes in 15,306 BS cows using additive and non-additive models. Ten additive QTL surpassed our significance threshold of p < 5e-08. Non-additive association testing revealed two QTL that remained undetected with the additive model. Two pronounced association signals on BTA25 caught our attention: an additive QTL that had been identified previously in BS cattle by Guo et al. (55) with the top associated SNP at 1,270,205 bp (b = 0.864, p = 1.205e-37); and a non-additive QTL, with the top associated SNP at 14,535,327 bp (d = 0.750, p = 2.35e-21) (Fig. 3A). To the best of our knowledge, the non-additive QTL had not been reported before in cattle. Distribution of stature measurements in individuals grouped by their genotypes at the top associated SNP from the non-additive GWAS confirmed recessive inheritance of the alternate allele (Fig. 3B).

**Figure 3.**
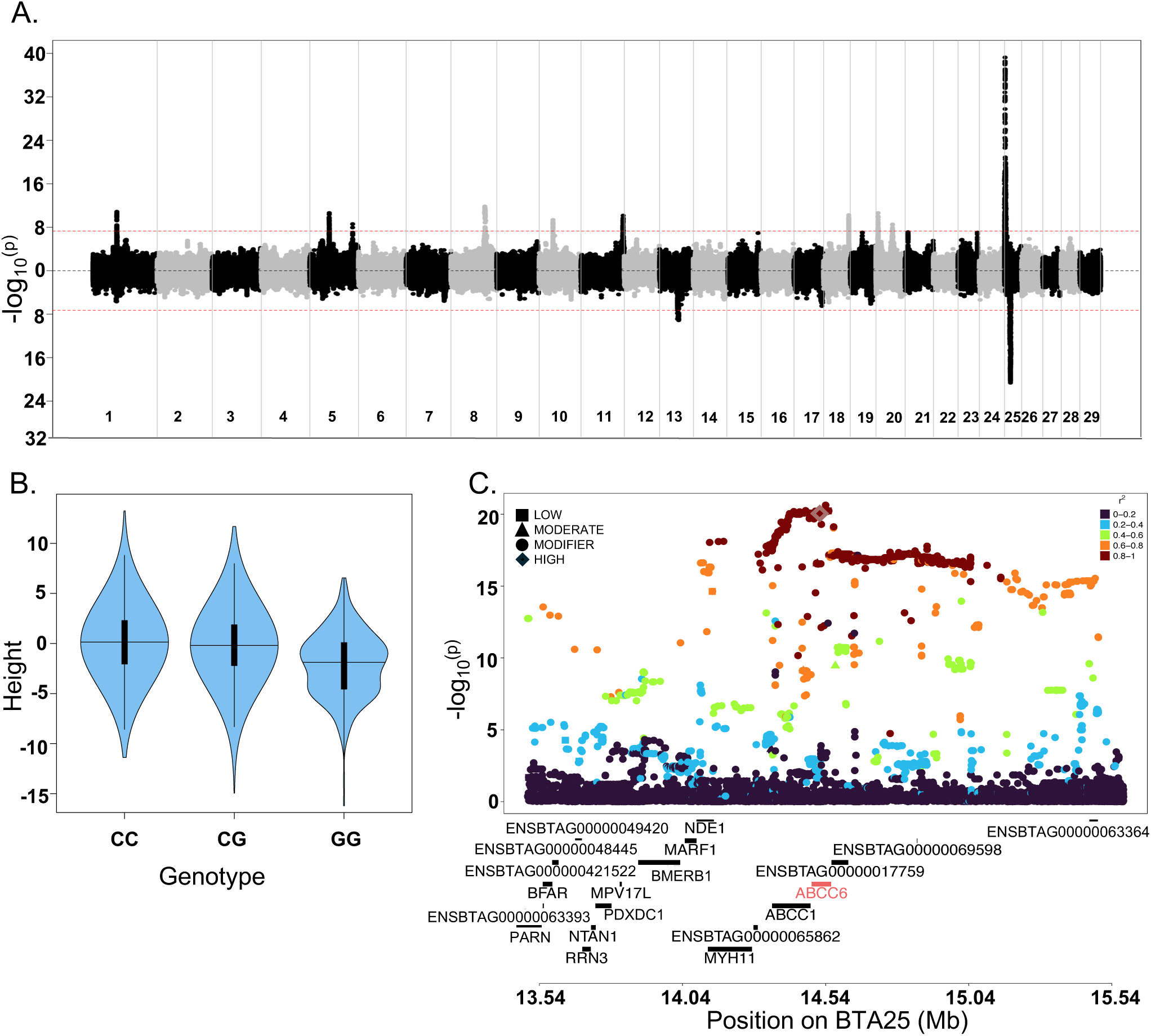
Genome-wide association testing between imputed sequence variants and statures in 15,306 Brown Swiss cows. **A.** Mirrored Manhattan plot showing association test results from the additive (top) and non-additive model (bottom). The red horizontal line indicates the genome-wide significance threshold (5e-08). **B**. Violin plot showing the distribution of statures in samples grouped by the genotypes at the top associated SNP (14,535,327 bp) at the non-additive QTL on BTA25 (C is the reference allele, and G is the alternative allele). **C.** Zoom plot of a 2-Mb region centered on the top variant at the non-additive QTL on BTA25. SNPs are coloured by the strength of LD (r^2^) with the top associated SNP. The high-impact splice variant in *ABCC6* (red) in high LD with the top associated SNP is highlighted. VEP-predicted SNP effects are indicated using different shapes.

To quantify the effect of the non-additive QTL on BTA25, we regressed stature on the genotypes of the top associated SNP which were coded as 1 for homozygotes for the alternate allele and 0 for all other animals. This model, which also included age and days since calving as fixed effects and expert and farm as random effects, estimated a significant decrease of 2 cm in status (beta = –1.97, p =1.72e-29) for individuals homozygous for the alternative allele (G|G) compared to heterozygotes (C|G) or individuals homozygous for the reference allele (C|C). Three quarters (266/354; 75%) of the cows that were homozygous for the alternative G allele also carried a ROH (medium-sized in 74% cases) encompassing the top associated SNP (Additional file 2, Supplementary Table S1), with an average ROH length of 1727 kb (± 5183kb). While only 40% of cows homozygous for the reference C allele carried a ROH encompassing the top associated SNP (Additional file 2, Supplementary Table S1). The genotype frequencies at this variant did not deviate from the Hardy-Weinberg proportions (p = 0.90, MAF = 0.15).

Another non-additive QTL was located on BTA13 (d = 1.93, p = 7.43e-10), with the top associated SNP at 47,076,701 bp. The genotype frequencies at the top associated SNP showed deviation (p = 2.00e-07) from Hardy-Weinberg proportions primarily resulting from an underrepresentation of alternative homozygous genotypes (18 observed vs. 49 expected).

### A splice donor variant of *ABCC6* is a candidate causal variant for the non-additive QTL on BTA25

The pronounced non-additive QTL for stature on BTA25 prompted a detailed investigation. We focused on a 2-Mb window spanning from 13,535,327 bp to 15,535,327 bp centered on the most significantly associated SNP (rs211632402 located at 14,535,327 bp, p = 2.35e-21), which was an intronic variant in *ABCC6* encoding ATP binding cassette subfamily C member 6. This region encompassed 20 protein-coding genes and 24,102 variants, including 1,645 with p values below 5e-08. The p values of all variants outside this window were at least six orders of magnitude higher than that of the top variant.

Within the 2-Mb region, 3,194 variants were in high LD (r² > 0.8) with rs211632402. Variant effect predictions indicated that nine of these variants had a moderate or high impact on protein function (Additional file 3, Supplementary Table S2). However, rs447836030, located at Chr25:14,515,474 and overlapping a canonical splice donor site of the 18^th^ *ABCC6* intron, was the only variant predicted to have a high impact on the resulting protein’s function. This variant had a p value of 8.96e-21, which is slightly less significant than that of the top variant.

ATP Binding Cassette Subfamily C Member 6 plays an important role in cholesterol transport and lipid metabolism (56). Cholesterol and lipid metabolism abnormalities have been shown to correlate negatively with height (57). The dysfunction of *ABCC6* is associated with various disorders such as Pseudoxanthoma elasticum in humans, a progressive metabolic disorder characterized by mineralization of the skin and elastic tissues (58). Mice deficient in ABCC6 have multiple partly age-dependent abnormalities of their vertebral bones (59).

Transcriptomic data from the Cattle Genotype-Tissue-Expression project indicate that *ABCC6* is predominantly expressed in the liver where transcript levels exceed 30 transcripts per million (TPM) (60). However, putative impacts of the splice donor variant on gene expression and splicing have not been investigated so far because rs447836030 segregates primarily in BS cattle for which comprehensive liver transcriptome data are not available. Analysis of a previously established cohort of 117 deep RNA and DNA sequenced BS bulls (39) revealed that *ABCC6* is also expressed in testis, albeit at a low level (1.97 ± 0.4 TPM). The rs447836030 T allele, which is predicted to abolish the splice donor site (GT => CT; noting that *ABCC6* is transcribed from the reverse strand), was observed in the heterozygous state in 30 bulls and in the homozygous state in 2 bulls, corresponding to an allele frequency of 14.5%. *ABCC6* mRNA expression is significantly (p = 8.18e-06) lower in animals carrying the rs447836030 T allele (Fig. 4A). Analysis of exon-specific expression suggests that the rs447836030 T allele causes the skipping of *ABCC6* exon 18 (ENSBTAE00000164650_14515642_14515475), which is predicted to result in an in-frame loss of 56 amino acids (Fig. 4B).

**Figure 4.**
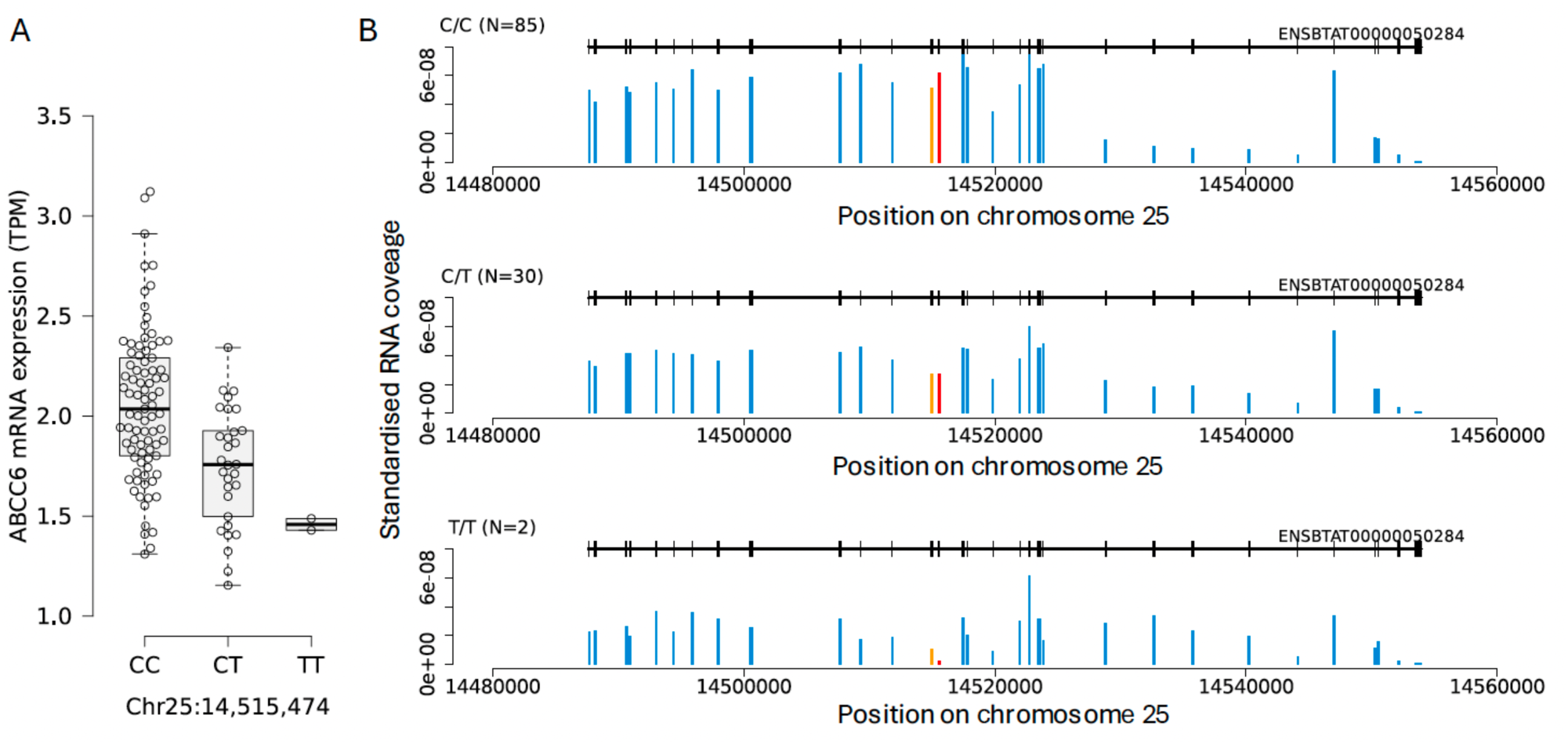
***ABCC6* mRNA expression in testis tissue**. A) Boxplots representing the mRNA expression of *ABCC6* in 117 mature bulls. B) Exon-specific expression of ABCC6 in the same animals. The upper, middle, and lower panels represent values averaged over 85, 30, and 2 bulls, respectively, that are homozygous for the reference allele (0/0), heterozygous (0/1), and homozygous for the alternate allele (1/1) of rs447836030. The black colour indicates the exon-intron structure of *ABCC6* transcript ENSBTAT00000050284. Blue bars represent the average standardised exon coverage in testis tissue. The red and orange colour indicates the 18^th^ and 19^th^ exons of *ABCC6*.

Differential splicing analysis revealed that a substantial fraction of *ABCC6* mRNA transcripts in animals carrying the rs447836030 T allele also lack exon 19 (ENSBTAE00000164654_14515023_14514849) (Additional file 4, Supplementary Figure S2), which explains the relatively low expression of exon 19 revealed in the exon-specific expression analysis. The skipping of both exons is predicted to result in a premature termination codon at amino acid position 778. The overall low abundance of *ABCC6* mRNA in animals carrying the rs447836030 T allele compared to animals homozygous for the reference allele possibly suggests that the severely truncated *ABCC6* transcript is subjected to nonsense-mediated mRNA decay.

### ROH are enriched for deleterious variants

We studied the distribution of deleterious and non-deleterious variants within ROH to investigate if certain types of ROH are enriched for high-impact variants. The numbers of homozygous non-deleterious (5252 ± 192) and deleterious (878 ± 56) alleles per genome were, as expected (17,36,61), positively correlated with F_ROH_ with values of r_n_= 0.718 (p < e-22) and r_d_ = 0.652 (p < e-22) respectively. Positive correlations (r_n_ > 0.305, r_d_ > 0.351, p < e-22) were also observed for short, medium and long ROH (Additional file 5, Supplementary Figure S3). Model M1, regressing the proportions of non-deleterious (baseline) and deleterious allele homozygosity inside ROH on F_ROH_ (see methods) showed a significantly higher intercept (β2 = 0.058, p = 0.55e-16) and higher slope (β3 = 0.078,p = 1.5e-07) than baseline, indicating that the proportion of homozygosity for deleterious alleles inside ROH consistently exceeds that of non-deleterious alleles at increasing values of F_ROH_ (Additional file 5, Supplementary Figure S3A). For instance, a highly inbred cow with a genomic inbreeding coefficient of 0.49 carried ∼ 43% more (1036 vs 726) deleterious alleles in homozygous form compared to a cow with a genomic inbreeding coefficient of 0.19. This increase was considerably less (23.5%; 5794 vs 4690) for the non-deleterious allele homozygosity.

The same regression model, regressing the proportions of non-deleterious (baseline) and deleterious allele homozygosity inside short, medium and long ROH, respectively, on F_short_, F_medium_ and F_long_ demonstrated that the slope for the proportion of deleterious allele homozygosity within ROH of all size categories was significantly higher than that of non-deleterious allele homozygosity (p < 0.05) (Additional file 5, Supplementary Figure S3B-D).

Confirming the stronger enrichment of deleterious allele homozygosity inside ROH compared to non-deleterious allele homozygosity, we sought to investigate the differences in enrichment across ROH of different size categories using model M2. The model was fit three times to compare the proportion of deleterious allele homozygosity inside (i) short vs medium, (ii) short vs long and (iii) medium vs long ROH (see Methods), defining the longer of the two ROH categories as the baseline. Short ROH had a significantly higher slope than long (β_3_ = 0.259, p = 5.06e-03) and medium ROH (β_3_ = 0.910, p = 3.94e-29), suggesting that the increase in the proportion of deleterious allele homozygosity in short ROH is higher per unit increase in F_short_ compared to this increase in long and medium ROH per unit increase in F_long_ and F_medium_ respectively. Between the medium and long ROH, the increase in the proportion of deleterious allele homozygosity was lower for medium ROH (β_3_ = –0.651, p = 3.33e-192) (Additional file 6, Supplementary Figure S4).

## Discussion

Our study provides evidence for inbreeding depression (ID) on stature in a large cattle population. A one-percent increase in the genomic inbreeding coefficient (F_ROH_) was associated with a 0.076 cm decrease in stature measured as height at the sacral bone, equivalent to 0.0195 standard deviations of the trait. This magnitude of ID on stature is comparable to estimates in other cattle breeds such as Holstein and Belgian Blue (0.0097 and 0.0234 standard deviations) (62–65) and to ID estimates in humans (between 0.029 and 0.037 standard deviations) (66,67). These findings suggest that the impact of inbreeding on stature may be similar across these species.

ROH of all size categories were associated with ID on stature in our data. However, we could not conclusively determine if the long ROH of recent origin or the short ROH of more ancient origin contributed more to this ID. The long ROH had the most significant effect. In contrast, the short ROH had the largest effect, which remained significant but was accompanied by a large standard error. It is possible that our study was underpowered to accurately estimate the effects of short ROH, which are expected to be shared across individuals and, hence, less variable (62).

To gain further insights into the possible adverse effects of inbreeding on fitness, we investigated the distribution of deleterious and non-deleterious alleles in ROH. This analysis showed that ROH were enriched for deleterious alleles, suggesting that the accumulation of deleterious alleles in homozygous form might underpin the observed adverse effects of inbreeding. The accumulation rate of deleterious allele homozygosity was the strongest within short ROH. This aligns with findings in other cattle breeds (18,68) but contradicts observations from human studies (17). While the lowest rate of accumulation of deleterious allele homozygosity was observed for medium ROH in our data, a previous study involving three other breeds suggested that deleterious allele homozygosity accumulated lowest in long ROH (18). Different patterns observed across breeds may have resulted from distinct genetic events, such as varying selection pressure and directions, shaping patterns of deleterious allele homozygosity across breeds (22). However, it is also possible that using imputed genotypes may have affected ROH classification. Some of the short ROH are possibly fragments of longer ROH which were broken due to imputation or genotyping errors. Nevertheless, the disproportionate (stronger) enrichment of ROH for deleterious allele homozygosity was consistent across ROH size categories. Increased burden of deleterious allele homozygosity at increased inbreeding underlines the necessity for proper management of inbreeding. The simultaneous presence of accumulated deleterious alleles (that individually may have milder effects) in ROH, particularly if their effects are synergistic, can pose a strong risk of disease phenotype (17).

Our 8^2^ estimate for stature of 3.48% aligns well with values previously estimated (4.9-6.9%) (29,69) in other cattle breeds. Identifying a recessive QTL on BTA25, which contributes disproportionately to the 8^2^ of stature, shows that non-additive GWAS can reveal loci contributing to dominance heritability and possibly ID. However, a much larger sample size might be necessary to identify more non-additive loci, as only one relatively common variant with a recessive effect was detectable in our data.

Stature and other easily measurable traits can serve as proxy phenotypes to uncover variants associated with hitherto undetected recessive disorders in the absence of formal disease classifications through non-additive association testing (29). Using this approach, we identified a novel recessive QTL for stature on BTA25. The effect size of this QTL is comparable to that of some recessive QTL for stature reported by Reynolds et al. (29) which were also associated with diseases. While Reynolds et al. observed homozygosity depletion in 4 out of 7 recessive QTL, we found no deviation from Hardy-Weinberg proportions for the top associated SNP at the non-additive BTA25 QTL. Moreover, all animals considered in our study were cows which had at least one offspring. This suggests that the QTL primarily contributes to quantitative variation in stature rather than causing a disease with a secondary effect on growth. However, since all the animals considered in our study were phenotyped within the first lactation, we cannot rule out the possibility of a late-onset disorder. Our findings add to existing literature on the pronounced effect of individual non-additive loci on complex traits in cattle (70,71).

A splice donor variant within *ABCC6* exhibits high LD with the non-additive QTL. Transcriptomic data supports this variant as a strong candidate for causality, as it induces exon skipping. This variant has two consequences; the skipping of exon 18 results in a protein that lacks 56 amino acids, and the skipping of exons 18 and 19 likely renders the resulting protein non-functional. The functional consequences of this splice donor variant were observed in testis tissue and need to be validated in other tissues, particularly in the liver and kidney, which are the primary sites of *ABCC6* expression. Given that variants affecting orthologs of bovine *ABCC6* had been shown to cause different disorders in other mammals (58,59), a thorough phenotypic examination of cows homozygous for the splice site variant in LD with the QTL is warranted at different life stages to investigate if other traits are impacted as well.

Most deleterious variants tend to be recessive and accumulate at low frequencies (72). This implies that detecting recessive effects in genetic studies requires a substantially larger sample size compared to detecting additive effects (73). A recessive allele identified in our study segregates at a frequency of 15% in BS cattle. A five-times larger cohort facilitated the identification of rarer non-additive variants with frequencies as low as 0.1% in Holstein cattle (29). This suggests that the sample size of our study is too small to detect rare recessive variants contributing to the genetic architecture of ID. An ever-increasing number of genotyped cattle with excellent phenotypic records will allow powerful studies to uncover both additive and non-additive variants contributing to the genetic architecture of complex traits and ID. Cattle and humans have distinct demographic histories and vastly different effective population sizes. A much smaller number of recessive variants are expected to segregate in cattle populations but at higher frequencies (26). This makes cattle a valuable model organism for studying non-additive genetic architectures as biobank-scale cohorts with both genotypes and phenotypes are available (74).

## Conclusions

This study provides evidence of inbreeding depression on stature in BS cattle, driven by ROH of different size categories. A non-additive GWAS revealed a novel deleterious recessive QTL, with homozygotes for its top-associated SNP predominantly located within ROH, contributing to inbreeding depression. A candidate causal variant was subsequently identified in the *ABCC6* gene. Further research is required to investigate whether dysfunction of this gene has detrimental effects in the later life stages of cattle.

## Declarations

### Ethics approval and consent to participate

Not applicable

### Consent for publication

Not applicable

### Availability of data and materials

Genotype and phenotype data analyzed in this study are owned by Braunvieh Schweiz and not publicly available. Summary data from the GWAS and ROH are available from the corresponding author upon reasonable request.

## Authors’ contributions

Preparation of genotype and phenotype data: QH NKK HP FS; GWAS, heritability estimation, assessment of inbreeding depression, analysis of ROH: QH NKK HP; QTL fine mapping and interpretation of results: QH NKK HP JD; Manuscript writing and revision: QH NKK HP. All authors read and approved the final manuscript.

## List of abbreviations

ID: Inbreeding depression
ROH: Runs of homozygosity
BS: Brown Swiss
OB: Original Braunvieh
GWAS: Genome-wide association study
QTL: Quantitative Trait Loci
SNP: Single nucleotide polymorphism
LD: Linkage disequilibrium

## Acknowledgements

We thank Braunvieh Schweiz for providing pedigree, phenotype and genotype data of BS cattle.

## Supplementary Information

**Additional file 1 Supplementary Figure S1: Scatter plot of the first two Principal Components used for breed identification**

PCA was performed using autosomal SNP genotypes. Samples without phenotype data are plotted in grey. The blue points correspond to BS (n=15,306), the red points represent OB (n=3,325), and the black points indicate mixed breeds(n=277).

**Additional file 2 Supplementary Table S1: Genotype distribution of the top associated SNP at the BTA25 recessive QTL with respect to ROH**

Genotype distribution (C|C, C|G, and G|G) of the top associated SNP (rs211632402) at the non-additive QTL within and outside ROH.

**Additional file 3 Supplementary Table S2: List of candidate causal variants for the non-additive QTL on BTA25**

Variants in high LD (>0.8) with the top associated SNP (s211632402) and predicted to have HIGH or MODERATE impact.

**Additional file 4 Supplementary figure S2: differential splicing analysis on *ABCC6***

Differential splicing of bovine *ABCC6* in testis. A) Partial structure of bovine *ABCC6*. The black boxes represent exons 16 – 20. Differently coloured lines are 6 splice junctions of splicing cluster 57171 identified by Leafcutter. B) Percent spliced in (PSI) values of 117 bulls. The different colours refer to the splice junctions as in A). Red colour indicates the junctions that support the skipping of exons 18 and 19 (ENSBTAE00000164650_14515642_14515475 and ENSBTAE00000164654_14515023_14514849).

**Additional file 5 Supplementary Figure S3: Proportion of deleterious and non-deleterious allele homozygosity within ROH**

Regression of proportion of deleterious allele homozygosity and non-deleterious allele homozygosity on the proportion of genome in A. ROH of any size (F_ROH_). B. short ROH (F_short_). C. medium ROH (F_medium_). D. long ROH(F_long_) in 15,306 individuals. The Pearson correlation coefficient between the deleterious (r_d_) and non-deleterious (r_n_) allele homozygosity and ROH coverage are indicated at the top left corner. The intercepts (β_2_) and slopes (β_3_) from the regression model M1, along with p values (one-tailed t-test), are also indicated.

**Additional file 6 Supplementary Figure S4: Proportion of deleterious allele homozygosity in short, medium and long ROH**

Regression of proportion of deleterious allele homozygosity versus proportion of genome in ROH of different size categories. The intercepts (β2) and slopes (β3) from the regression model M2, along with p values (one-tailed t-test), are indicated in the legend.

