## Supplementary figures and images for "Evidence of inbreeding depression on stature in Brown Swiss cattle"

### Supplementary Figure S1

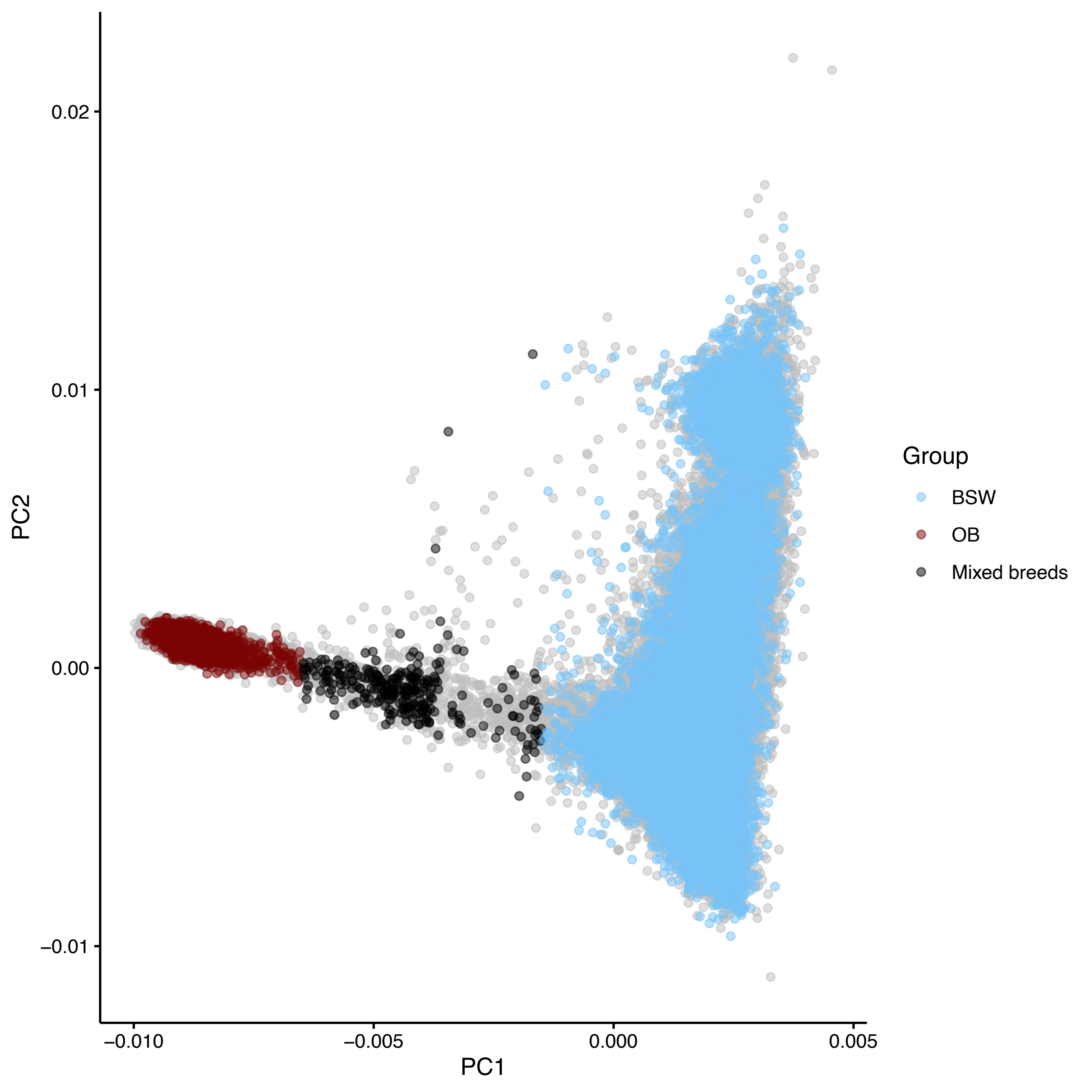

### Supplementary Figure S2

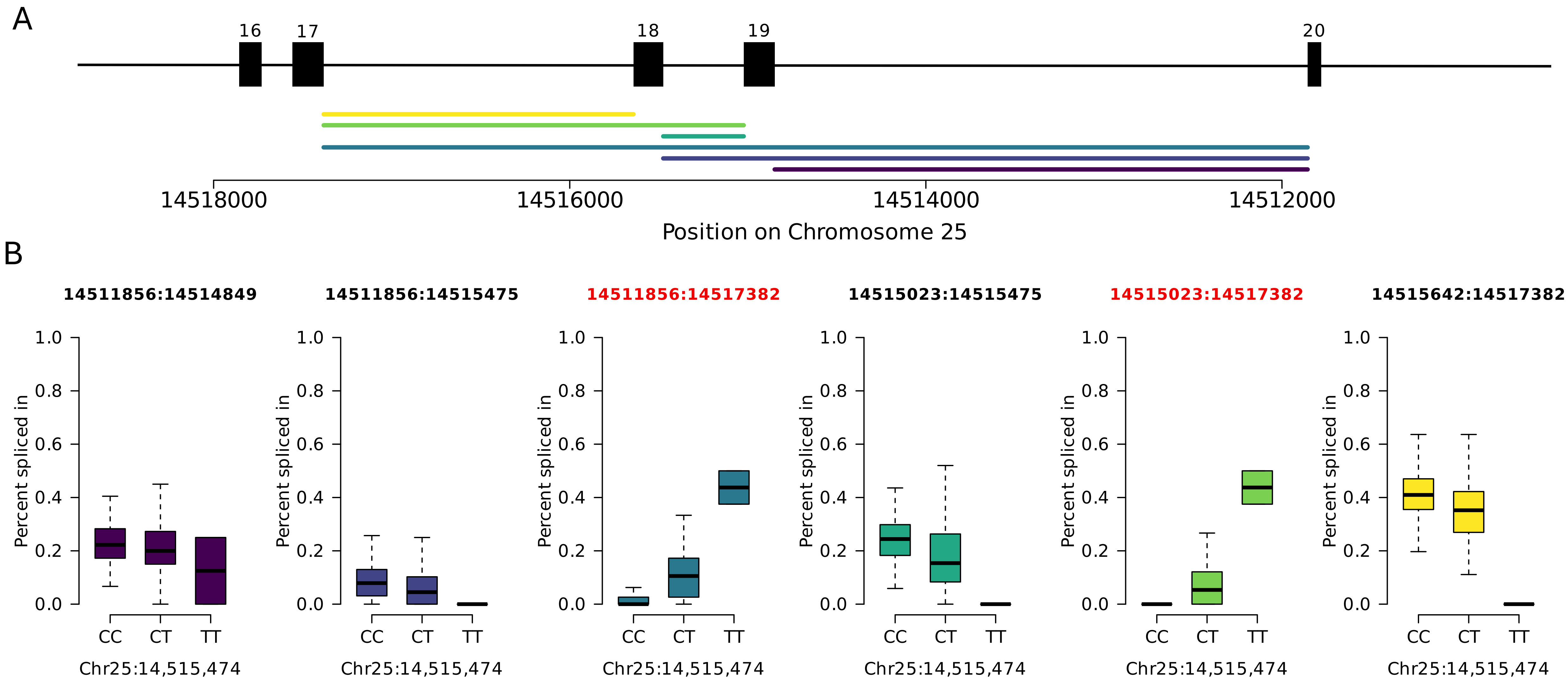

### Supplementary Figure S3

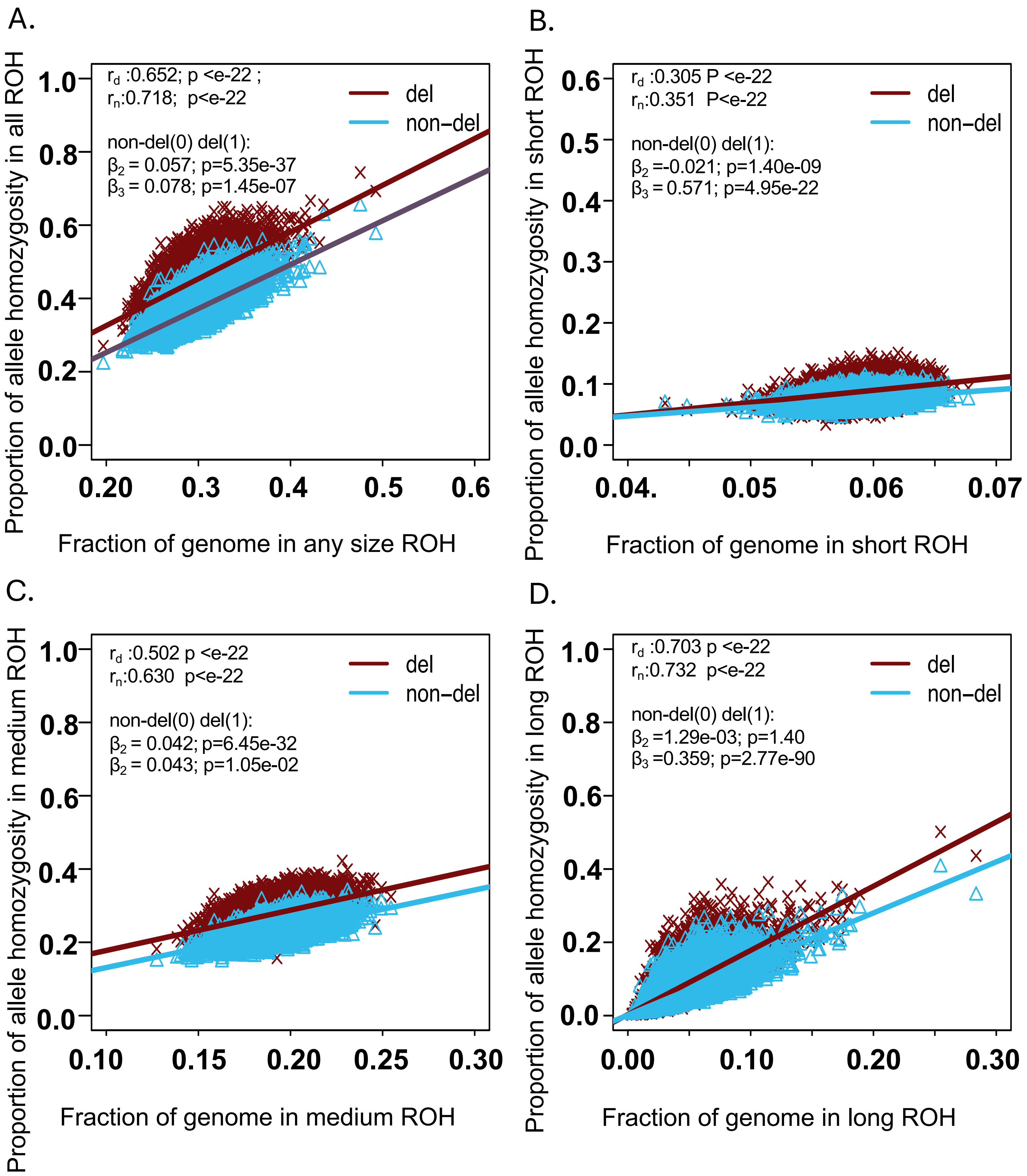

### Supplementary Figure S4

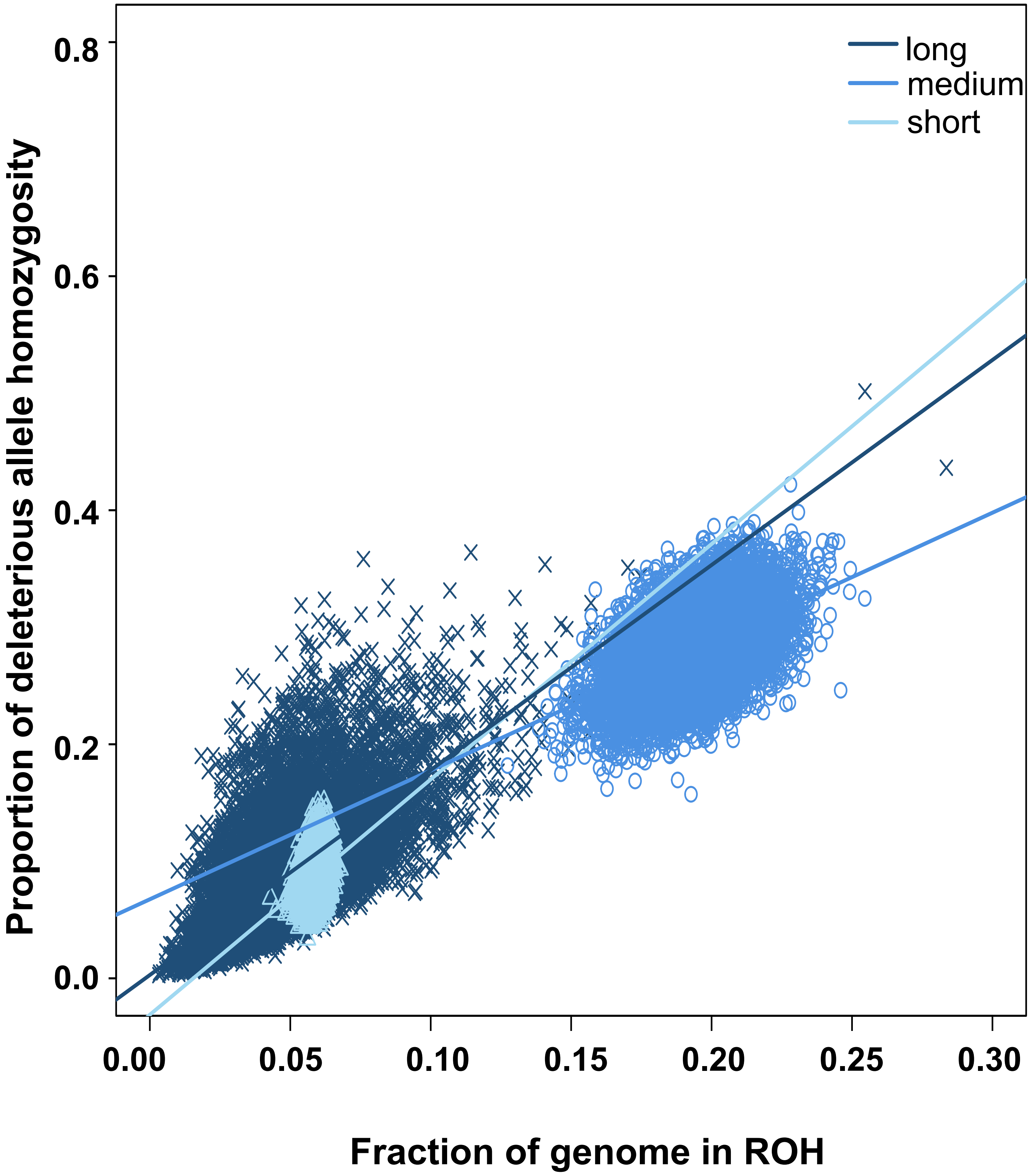


long (0) vs short (1) : $\beta_2$ = -0.033 p = 5.8e-09 ; $\beta_3$ = 0.259 p = 5.06e-03

long (0) vs medium (1) : $\beta_2$ = 0.064 p = 2.13e-82 ; $\beta_3$ = -0.651 p = 3.3e-192

medium (0) vs short (1) : $\beta_2$ = -0.098 p = 7.38e-83 ; $\beta_3$ = 0.910 p = 3.94e-29
